# Visualization of lake succession as indicated by macrophyte species composition and trophic levels in lakes subdivided by volcanic eruption at Akan Caldera, Hokkaido, Japan

**DOI:** 10.1101/2021.05.05.442726

**Authors:** Isamu Wakana, Yasuro Kadono, Jotaro Urabe, Yuki Tamura, Yoshifusa Suzuki, Hiroyuki Yamada, Yoichi Oyama, Keiji Wada, Takeshi Hasegawa, Masashi Ohara

## Abstract

It is not feasible to continuously observe ecological succession in lakes because of the long time-scales generally involved. Thus, the process has been inductively deduced by comparing many lakes with different succession states, or indirectly simulated by tracking studies of smaller water bodies, experiments using microcosms or mesocosms, and reconstruction of lake history by sediment analysis. However, the reality of succession processes in large lakes with slow succession is not well understood, and new approaches are needed. Theoretically, in a group of large and small lakes of similar ages and with similar initial and watershed environments, the rate of nutrient accumulation in each lake depends on the ratio of watershed area to lake size, and the lakes are predicted to evolve to different trophic levels over time. Here, we tested this hypothesis on the 10 lakes of varying sizes in Akan Caldera, Japan, which were formed thousands of years ago by fragmentation due to volcanic eruptions within the caldera. Topographic and water quality studies showed that the ratio of accumulated watershed area to lake size (area and volume), expressed logarithmically, had a positive linear regression with the total phosphorus concentration, an indicator of trophic level. The trophic levels of the lakes were diverse, including oligotrophic, mesotrophic, and eutrophic types in the traditional “lake type” classification based on total phosphorus and chlorophyll-a concentrations. Furthermore, 21 species of aquatic macrophytes were observed by a diving survey, and the plant species composition was classified into five groups corresponding to the trophic status of the lakes, indicating a conventional “hydrarch succession”. The diversity of water quality and aquatic vegetation in a group of lakes with similar origins paves the way for new comparative studies of lakes, including large lakes.

## Introduction

Ecological succession within lakes is not only a major component of freshwater ecology and limnology, but also a platform for diagnosis and management of worldwide deteriorating aquatic environments resulting from human activity since the 20th century [1–7]. In general, lake succession proceeds thusly: when a lake is first formed the water is oligotrophic with a paucity of biota, but the trophic level subsequently increases, as trophic substances inflow from the watershed, and the lake basin shallows from accumulation of sediments. The diversity of biota and biomass increase or fluctuate throughout this process, and eventually the lake attains the state of bog or marsh [1–4, 8]. Historically, Forel, the “father” of limnology [4], explained this transition as analogous to human ageing [9], and this concept has been widely accepted due to both intuitive and experiential evidence [1, 2, 4, 10]. Forel also pointed out that it is impossible to follow a normal lake’s evolution, because of the immense time-scales required for a lake to disappear by sedimentation [9]. Therefore, this general picture of long-term succession has been indirectly formed by comparing many lakes with different trophic conditions [10–16], follow-up survey of small-scale dams or reservoirs [2, 17], experimental microcosms and mesocosms [4, 18–22], and historical reconstruction of sedimentation processes [5, 8, 23]. Despite these efforts, however, the perception of ecological succession in large lakes still relies on many assumptions, due to their long ageing process [2, 18, 24–26].

Is there any other approach to visualize lake succession processes, including within large lakes? In principle, the rate of eutrophication, a major factor in ecological succession, is determined by lake size, watershed area and trophic condition of the watershed [4]. Therefore, if trophic conditions are equal among different watersheds, the eutrophication rates, i.e., the rates of nutrient increase, should vary depending on the ratio of the watershed area to the lake size in each lake. In addition, given a group of lakes with similar formation time and initial environment, the concentrations of nutrients are expected to change with a positive linear regression between the ratios of watershed area to lake size. As described, such lakes can be considered as a series that evolved into different trophic levels, providing a new approach to visualizing lake succession. In practice, however, no study has yet demonstrated this relationship, because nutrient loading from the watershed is generally influenced by indigenous variables, such as land form, soil, erosion, local climate (temperature and rainfall), vegetation, and land use (farmland, factory and urbanization) in addition to the time effect [1–4, 10]. We therefore offer insight into this problem by comparing a group of lakes within a large caldera. The lakes of Akan Caldera, Hokkaido, Japan, were formed by volcanic eruptions thousands of years ago, which divided a huge caldera lake into several lakes of varying sizes [27–29]. These lakes share a similar terrestrial environment within the caldera. In contrast, the trophic levels in these lakes are known to be diverse, ranging from oligotrophic to eutrophic [30–33], and may show a series of “lake successions” or “lake types” in the natural ecological succession of lakes [2, 5, 10]. In this paper, by studying the topography and water quality of Akan Caldera Lakes, we tested the hypothesis that lakes with a lower ratio of watershed area to lake size are oligotrophic, while lakes with a higher ratio are eutrophic.

Furthermore, the lake ecological succession is also displayed as “hydrarch succession” according to changes in aquatic vegetation [2, 5, 10, 34]. The macrophytes, which are major players in this process, are known to change their species composition in response to changes in trophic levels, especially in eutrophic lakes affected by anthropogenic activity [35–38], but the natural hydrarch succession in large lakes is not fully understood, as is the case with lake succession or lake type. Through preliminary surveys across the Akan Caldera Lakes, we have observed that each lake has a unique macrophyte species composition, so we also examined the prediction that the species composition may correspond to the trophic level.

## Materials and methods

### Study area

Akan Caldera is situated at the southern end of the Akan-Shiretoko Volcanic Chain, a volcanic region in eastern Hokkaido, Japan [29]. The outer shape of the caldera is oblong (24 km east-west × 13 km north-south) [29], and a “central cone”, Mt. O-akan, rises at the center of its inner basin (Fig 1). Within the caldera there are lakes and marshes of various sizes surrounding Mt. O-akan (Fig 1). Lake watersheds are isolated from external input by the caldera wall [39, 40], and the water systems, connected by rivers or underground flow [27, 40, 41], are roughly divided north-south and join at the southern foot of Mt. O-akan, where they then discharge through Akan River, the notch in the south caldera wall (Fig 1A).

**Fig 1.**
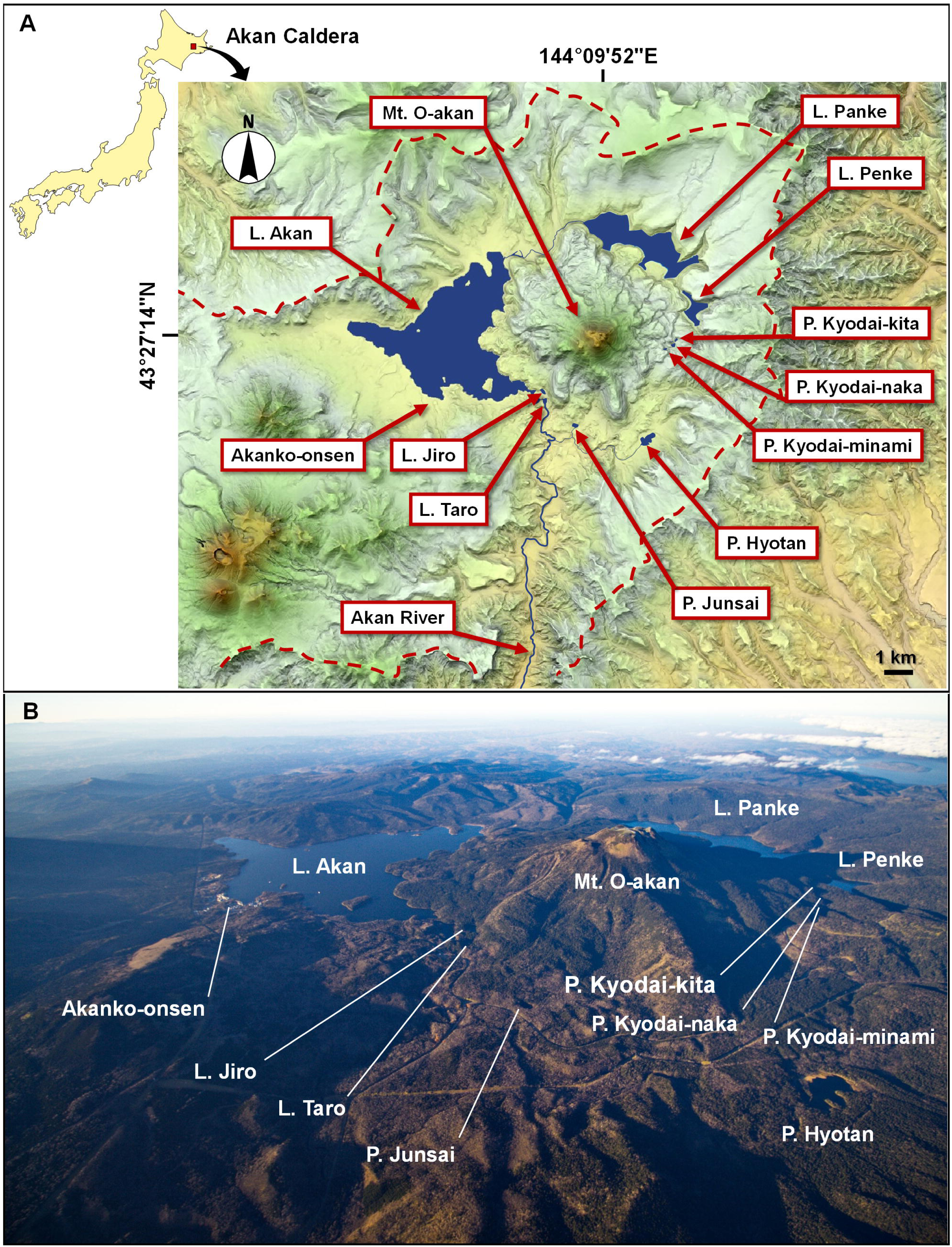
Map and landscape of Akan Caldera Lakes. (A) The watershed of Akan Caldera is isolated by a caldera wall shown by red dotted lines, but the western caldera wall is obscured by volcanos erupting after the formation of the caldera. Ten lakes surround Mt. O-akan within the caldera, and are connected by rivers and underground flow from water systems to the north and south. The water flows converge south of Mt. O-akan and discharge is through the Akan River. The map was generated by Kashmir 3C [42], a map software. (B) Land area is mostly covered with subarctic forests except for a town south of Lake Akan, Akanko-onsen.

Large lakes are distributed from west to northeast of Mt. O-akan and smaller lakes are localized from south to east (Fig 1A). This peculiar lake arrangement is a result of the formation history of the caldera and Mt. O-akan. The Akan region has witnessed more than ten large volcanic eruptions in the last 1.5 million years, and the present oblong-shaped caldera was formed by the largest eruption of 200,000 years ago [27, 29]. After the last large eruption (150,000 years ago), a huge lake, “Ko-akanko” (ancient Lake Akan), was generated in the caldera [27–29]. The lake was narrowed by post-caldera volcanic activity in the southwest part of the caldera, and by 110,000 years ago landform of the inside caldera wall was almost completed [27–29]. A minor eruption of Mt. O-akan occurred 13,000 years ago slightly southeast of the center of the caldera, and it stopped when the lava flow reached the caldera wall, so that Ko-akanko was separated into large and small basins [27–29].

The developmental history of Mt. O-akan also shows that Pond Hyotan and Pond Junsai of the southern water system were first divided from Ko-akanko, and other lakes of the northern water system were formed 5,000 to 2,500 years ago [27–29]. The depth charts of the large lakes, Akan, Panke and Penke, show the remains of valleys on the bottom of the inside caldera wall extending as far as the base of Mt. O-akan [27, 39, 41]. This suggests that the water level of Ko-akanko was extremely low or the basin was exposed by discharge through the notch before the eruption of Mt. O-akan [27]. Therefore, the lakes of the northern water system are thought to have formed by re-flooding after damming by Mt. O-akan, but the timing of lake formation and the developmental processes are not fully understood.

Akan Caldera occupies a part of Akan-Mashu National Park (designated in 1934), and is mostly covered with subarctic forests except for a town on the south side of Lake Akan, Akanko-onsen (Fig 1B) [30]. Only Lake Akan has been developed as a sightseeing area due to the presence of Marimo, *Aegagropila linnaei*, a ball-shaped green alga designated a Japanese natural treasure [30, 43]. Since the 1950s, the increase in tourism has resulted in eutrophication from sewage discharge. This continued until the 1980s when public sewage treatment service was provided [44].

### Topography of lakes and watersheds

Lake area and boundary length were calculated using ARCGIS10 (Esri Japan Co.) based on the 1/25,000 numerical map data of the Geographical Survey Institute, Japan. Land area includes island areas.

Land watershed area was computed using the DEM10m data of the Geographical Survey Institute. After altitude data were changed into raster (altitude grid), subtle undulations were removed by the Fill tool, and bearing azimuth of flow was computed by the Flow Direction tool (north; 64, northeast; 128, east; 1, southeast; 2, south; 4, southwest; 8, west; 16 and northwest; 32). Accumulation value (number of cells accumulated toward the direction of flow) computed by the Flow Accumulation tool was extracted at accumulation values more than 30000 (sl30000) and more than 200 (sl200) by the Reclass command, and each watershed (ws30000, ws200) was computed by being grouped by every feeder of sl30000 and sl200 by the Stream Link tool. Finally, the Ws raster was converted into the polygon, and Ws30000 and ws200 were manually divided as a watershed for each lake, observing the DEM.

Lake volume and mean depth were calculated based on lake charts. Depth sounding for chart drawing was performed in autumn of 2014 in seven lakes, excepting Lakes Akan, Panke and Penke which already have lake charts. The depth of the entire lake was uniformly measured by a GPS fish finder (Lowrance HDS-8, Navico) on a motor boat, and the depth-sounding data were converted into contour drawings by chart drawing software, Reefmaster (ReefMaster Software Ltd.). Elevation of the lake surface was obtained by the GNSS survey. Residence time was calculated by dividing lake volume by inflow obtained through multiplying annual rainfall at 1,200 mm [30] by the accumulated watershed area (see Statistical analysis, below).

### Water quality

Measurements of physical and chemical variables and collection of lake water were performed twice in ten lakes of Akan Caldera from October to November 2013 and in July 2014. Secchi depth, water temperature (Temp), electrical conductivity (EC) and pH were directly recorded using a Secchi disk and portable sensors at the center of each lake. Water was collected at the same point using 2 l polycarbonate bottles and taken immediately to the laboratory. Dissolved oxygen (DO) and chemical oxygen demand (COD) were measured by titration with standard sodium thiosulfate solution and potassium permanganate solution, respectively [45]. Total nitrogen (TN) and total phosphorus (TP) were measured by an auto analyzer (AACS-II, Bran+Luebbe Ltd.) [46]. Additionally, an aliquot of the water sample was filtered onto Whatman GF/F glass fiber filters, and suspended solids (SS) measured gravimetrically after drying at 110 °C for two hours [45]. Chlorophyll-a (Chl-a), concentrated onto a Whatman GF/F glass filter, was quantified with a spectrophotometer (UV-1600, Shimadzu Co. Ltd.) after extraction using methanol (100%) [47]. The filtrate was used for measuring dissolved organic carbon (DOC) with a TOC meter (TOC VC, Shimadzu Co. Ltd.) [48].

### Macrophyte survey

Macrophytes were surveyed in 10 lakes of Akan Caldera by SCUBA diving or snorkeling from October to November 2013 and in July 2014. The maximum depth surveyed was 18 m in Lake Panke.

### Statistical analysis

A total of 31 variables were analyzed (S1 Table). The lake topographic factors were 11 variables: elevation, boundary length, lake area (LA), shore line development, maximum depth, mean depth, lake volume (LV), residence time, land watershed area (LWA), total watershed area (TWA) which is the sum of LA and LWA, and accumulated watershed area (AWA) which is the sum of the TWA above a given lake. LA and LWA correspond to the aquatic and terrestrial watershed areas specific to individual lakes, respectively, and AWA indicates the total watershed area including the upstream portion. Six variables reflected inflow and outflow of materials [3]: LWA-LA ratio, LWA-LV ratio, TWA-LA ratio, TWA-LV ratio, AWA-LA ratio and AWA-LV ratio. Water chemistry parameters consisted of ten variables: Temp, pH, DO, EC, SS, Chl-a, DOC, COD, TN and TP, and the macrophyte communities were defined by four variables: number of submerged species, floating-leaved species, free-floating species and total species. Secchi depth was excluded from the analysis due to missing data (S1 Table).

To explore the relationship among topography, water quality and macrophyte community, a correlation matrix of these standardized data was made (S2 Table) and variables with strong correlation coefficients (|r| ≥ 0.7) were investigated in detail. In lakes where macrophytes occurred, the presence/absence of individual species was converted into 1/0 data, and a cluster analysis was conducted by the Ward method using the Euclid distance. The BellCurbe (Social Survey Research Information), an add-in program for Excel, was used for the analysis.

## Results and discussion

### Water quality and lake types

The water quality of the 10 lakes was diverse (S1 Table), and classified as oligotrophic (2 lakes), mesotrophic (5 lakes) and eutrophic (3 lakes) according to Chl-a and TP concentrations (Fig 2A) [7]. Among these lakes, Pond Junsai (eutrophic) had brownish water with high TN (0.438 mg l^−1^), DOC (7.1 mg l^−1^) and Chl-a concentrations (12.5 μg l^−1^) (S1 Table), indicating dystrophic properties [5, 21].

**Fig 2.**
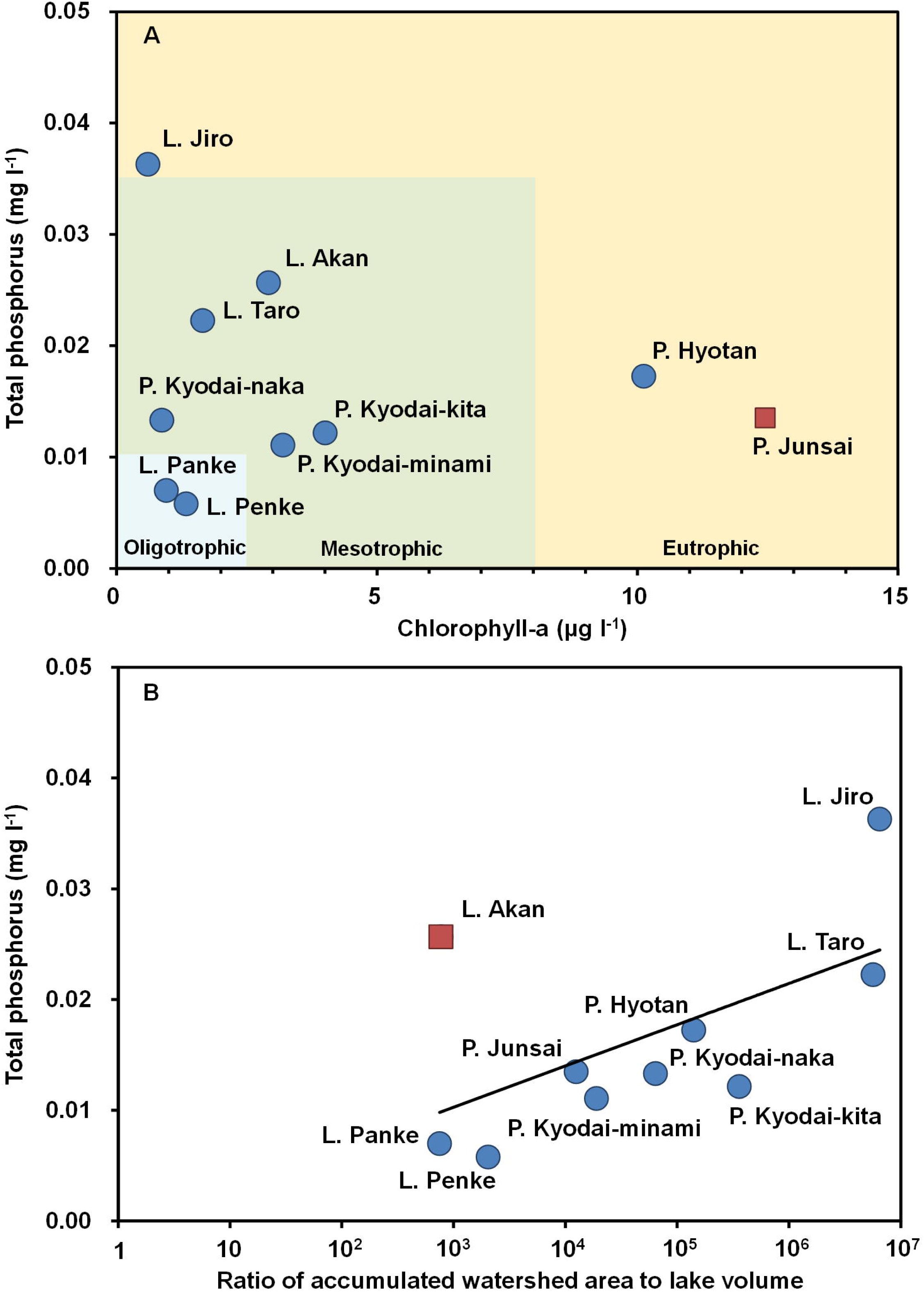
Aspects of trophic status in Akan Caldera Lakes. (A) A variety of lake types categorized by Chlorophyll-a and total phosphorous (TP) according to the Organization for Economic Co-Operation and Development (OECD) [7]. Even though the lakes share the same origin and most of them were formed at the same time, the lake types are diversified into oligotrophic, mesotrophic and eutrophic. Additionally, Pond Junsai (red square) has dystrophic characteristics (S1 Table). (B) Relationship between ratio of accumulated watershed area to lake volume (AWA/LV) and TP. The AWA/LV shows significant correlation with TP thought to be a parameter of eutrophication rate, but only Lake Akan (red square), subjected to anthropogenic eutrophication in the past, shows an unusually high TP.

Among ten water quality variables and 17 topographic characters, strong (|r| ≥ 0.7) and significant (*p* < 0.05, by test of no correlation) correlation coefficients were found between: TP and AWA, TP and AWA-LV ratio (AWA/LV), EC and AWA, EC and AWA-LA ratio (AWA/LA), EC and AWA/LV, pH and shoreline development, DO and AWA/LV, and Temp and elevation (Table 1). Although correlation coefficients between TP and AWA/LA, and DO and AWA/LA did not reach 0.7 and −0.7, respectively, both were statistically significant (Table 1).

**Table 1.**
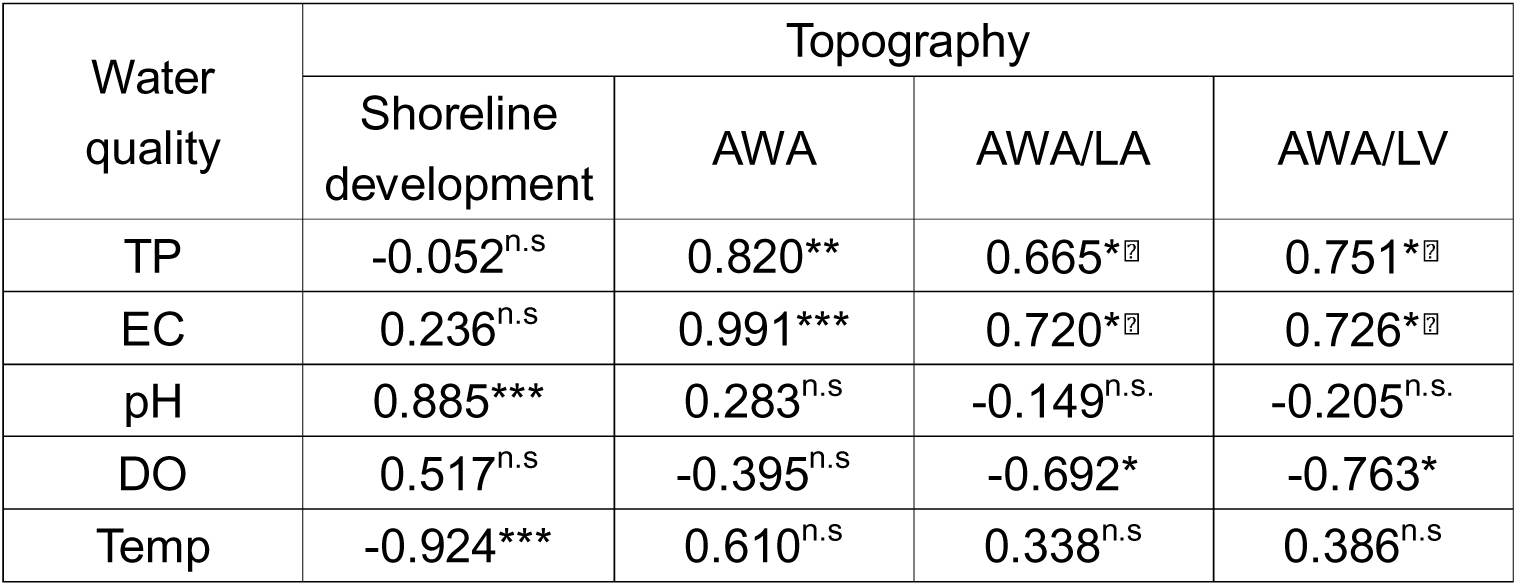
Combinations indicating strong correlation coefficients (r) between water quality variables and topographic characters.

| Water quality | Topography |  |  |  |
| --- | --- | --- | --- | --- |
|  | Shoreline development | AWA | AWA/LA | AWA/LV |
| TP | -0.052<sup>n.s</sup> | 0.820<sup>**</sup> | 0.665<sup>*☒</sup> | 0.751<sup>*☒</sup> |
| EC | 0.236<sup>n.s</sup> | 0.991<sup>***</sup> | 0.720<sup>*☒</sup> | 0.726<sup>*☒</sup> |
| pH | 0.885<sup>***</sup> | 0.283<sup>n.s</sup> | -0.149<sup>n.s.</sup> | -0.205<sup>n.s.</sup> |
| DO | 0.517<sup>n.s</sup> | -0.395<sup>n.s</sup> | -0.692<sup>*</sup> | -0.763<sup>*</sup> |
| Temp | -0.924<sup>***</sup> | 0.610<sup>n.s</sup> | 0.338<sup>n.s</sup> | 0.386<sup>n.s</sup> |

The original correlation matrix is available in S2 Table. TP: total phosphorus, EC: electrical conductivity, DO: dissolved oxygen, Temp: temperature, AWA: accumulated watershed area, AWA/LA: AWA-lake area ratio, AWA/LV: AWA-lake volume ratio, ***: *p* < 0.001, **: *p* <0.01, *: *p* < 0.05, n. s.: not significant, ⍰: Lake Akan as an outlier.

When two-dimensional plots were drawn on these 10 combinations, Lake Akan was discriminated as an outlier in TP and AWA/LA, TP and AWA/LV, EC and AWA/LA, and EC and AWA/LV (Table 1, Fig 2B). As previously noted, Lake Akan was subject to anthropogenic eutrophication in the second half of the 20th century [44]. We thus estimated phosphorus concentration before eutrophication. The oldest P_2_O_5_ data, 0.010 mg l^−1^, measured in Lake Akan in 1931 [49] was calculated at 0.004 mg l^−1^ phosphorus by the conversion formula which divides the value of P_2_O_5_ by 2.29 [10]. This result was close to the regression lines of TP and AWA/LA (r = 0.787, *p* < 0.05), and TP and AWA/LV (r = 0.881, *p* < 0.01) drawn for nine lakes except for Lake Akan (Fig 3A). The difference (0.022 mg l^−1^) from our observed data (0.026 mg l^−1^, S1 Table) is likely part of the increase from eutrophication, suggesting that before eutrophication, AWA/LA and AWA/LV of the lakes including Lake Akan had a tighter linear regression relationship with TP. Conversely, without including the outlier in the case of TP and AWA, the resulting plot when Lake Akan TP data were replaced by the above 0.004 mg l^−1^ drew away from the regression line (r = 0.795, *p* < 0.05, Fig 3B).

**Fig 3.**
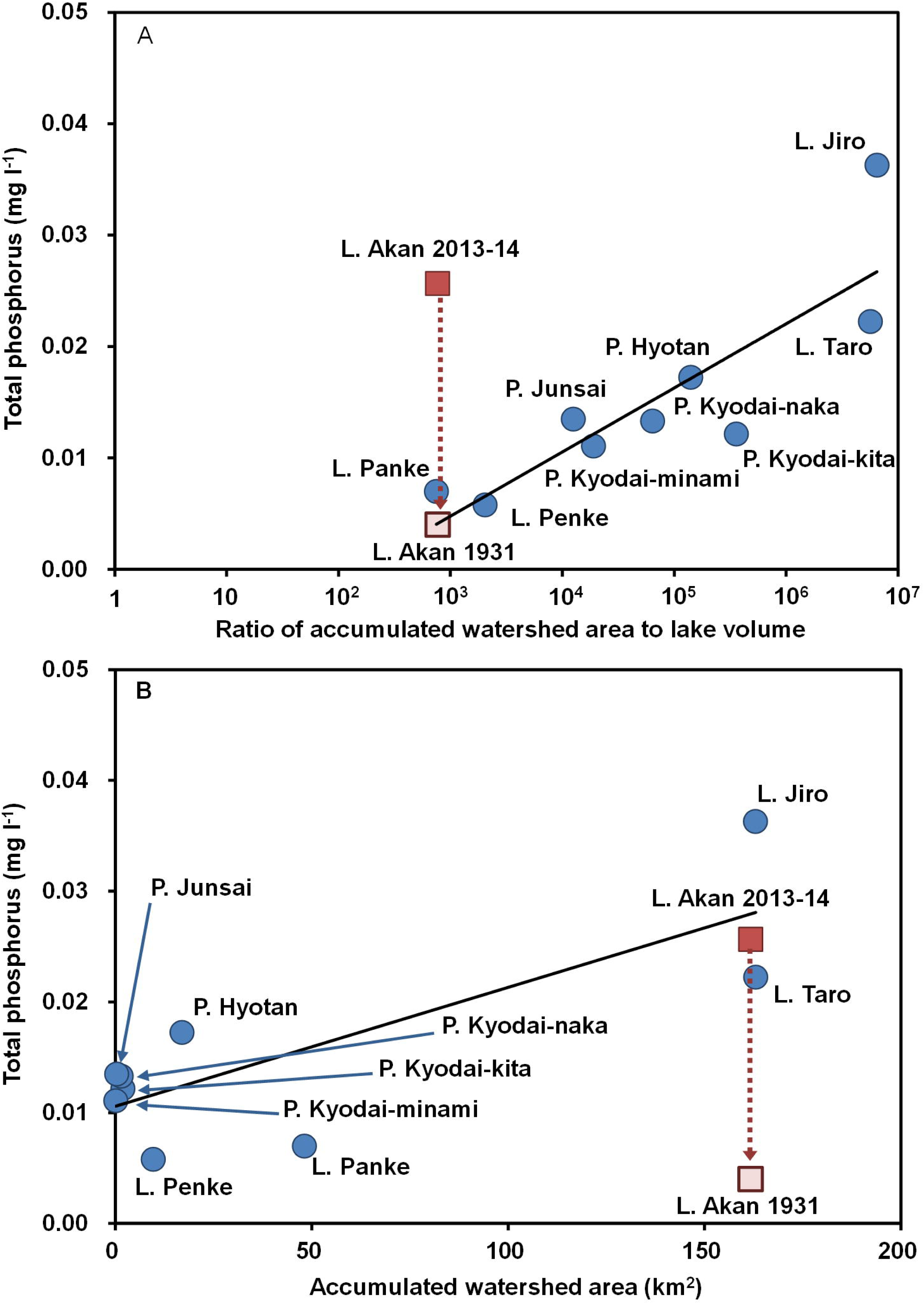
Estimation of phosphorus concentration in Lake Akan before eutrophication. Because Lake Akan was subject to anthropogenic eutrophication in the second half of the 20th century [44], phosphorus concentration before eutrophication was estimated based on data from 1931 [49]. Each regression line is drawn for nine lakes (blue circles) except for Lake Akan. In (A) relationship between ratio of accumulated watershed area (AWA) to lake volume and total phosphorus (TP), the 1931 estimate of phosphorus concentration for Lake Akan (light red square) is significantly lower than the 2013–14 TP data (dark red square), and is close to the regression line. Conversely, in (B) the relationship between AWA and TP, the same 1931 estimate is distant from the regression line.

A strong correlation was observed between EC and TP (r = 0.821, *p* < 0.01, S2 Table). The regression formula (*y* = 819.023*x* - 2.343) estimated the EC of Lake Akan at 0.933 mS m^−1^, near the regression lines of EC and AWA/LA (r = 0.973, *p* < 0.001,) and EC and AWA/LV (r = 0.979, *p* < 0.001) for the nine lakes when TP was 0.004 mg l^−1^ (Fig 4A). By contrast, the calculated result of Lake Akan EC drew away from the regression between EC and AWA without the outlier (r = 0.988, *p* < 0.001, Fig 4B). This suggests that the involvement of AWA may be invalid, along with the results of similar substitution of that between TP and AWA, shown in Fig 3B.

**Fig 4.**
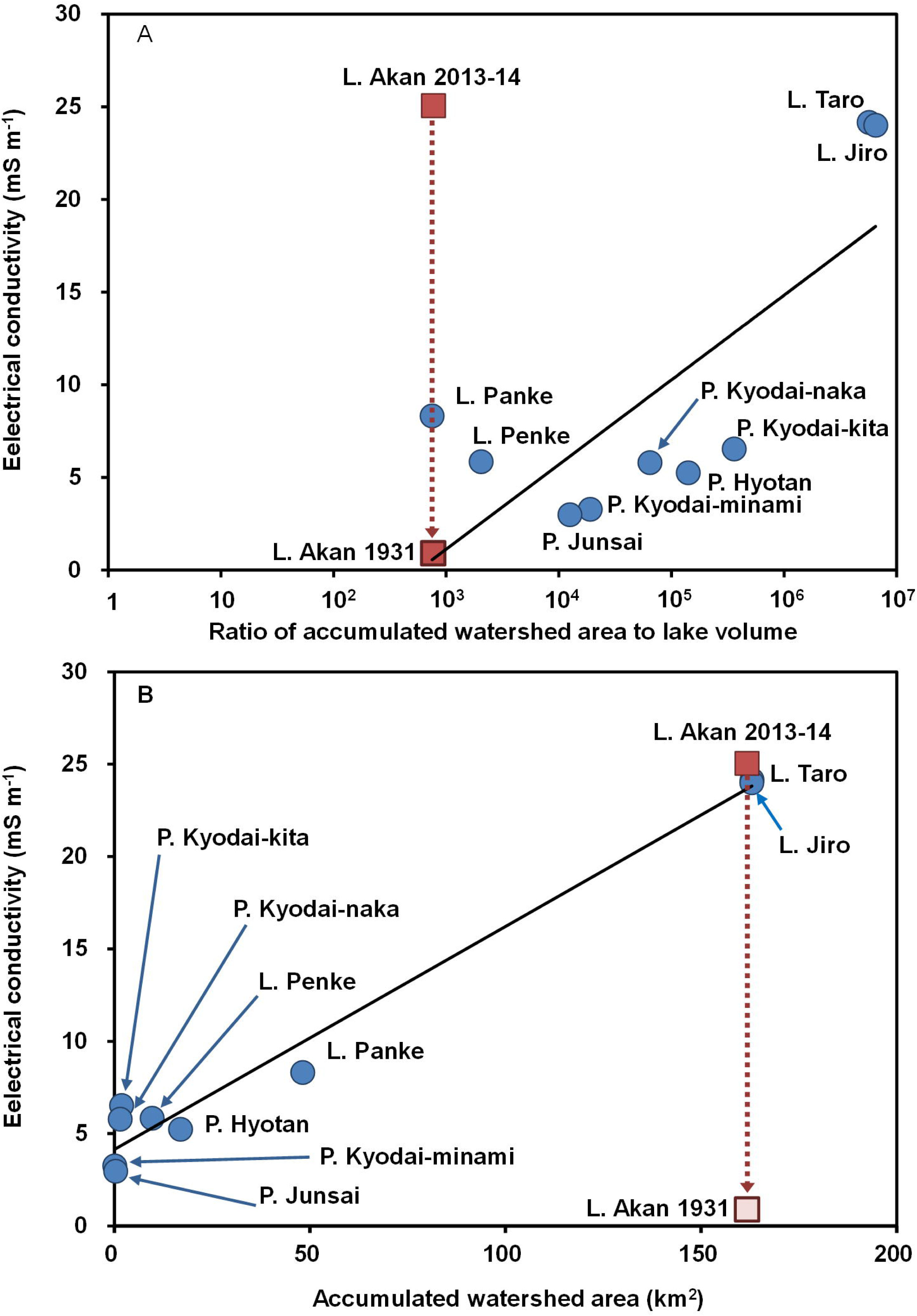
Estimation of electrical conductivity (EC) in Lake Akan before eutrophication. Because phosphorus concentration in Lake Akan was estimated to be significantly lower before eutrophication (Fig. 3), EC showing a high correlation with total phosphorus (TP) was also examined. The EC for 1931 was calculated at 0.933 mS m^−1^ by substituting the estimated phosphorus concentration for 1931 (0.004 mg l^−1^) into the regression formula for EC and TP. Each regression line in the graphs is drawn for nine lakes (blue circles) except for Lake Akan. In (A) relationship between ratio of accumulated watershed area (AWA) to lake volume and EC, the 1931 estimate of EC for Lake Akan (light red square) is significantly lower than the 2013–14 EC data (dark red square), and is close to the regression line, similar to the case of phosphorus in Fig. 3. Conversely, in (B) the relationship between AWA and EC, the same 1931 estimate is distant from the regression line, as in the case of phosphorus.

The regression line of pH and shoreline development was likewise drawn without including the outlier (Fig 5). Because shoreline development is proportional to length of the horizontal littoral zone, the correlation with pH was presumed to relate to consumption of CO_2_ through hydrophytic photosynthesis [2, 4, 50].

**Fig 5.**
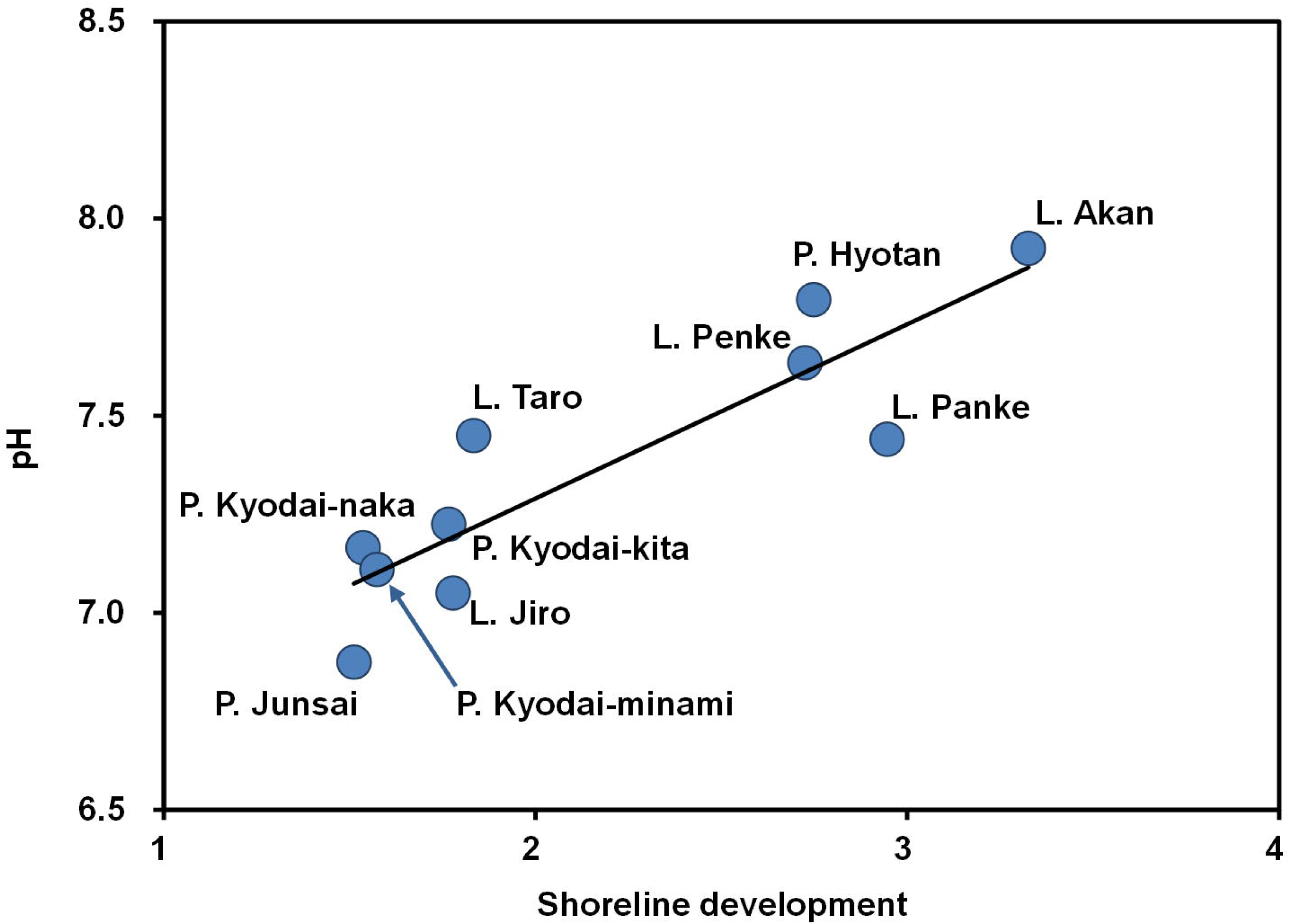
Relationship between shoreline development and pH in Akan Caldera Lakes. Despite the lakes having a rich diversity of lake basin sizes and water quality, including trophic levels (S1 Table), shoreline development, the degree of shoreline flexion, is strongly correlated with lake water pH.

The regression lines of DO and AWA/LA, and DO and AWA/LV were drawn with a slight negative slope due to the specific low DO data for Lake Jiro (Fig 6), and may not indicate an environmental gradient. Lake Jiro has no inflow and outflow rivers (Fig 1A), and the water appears to be supplied through underground flow from the upstream Lake Akan [51]. However, in addition to DO, Chl-a, DOC, and COD in Lake Jiro were the lowest concentrations among the Akan Caldera Lakes, and TP was the highest (S1 Table). Furthermore, a portion of the lake surface of Lake Jiro does not freeze in winter [51]. These results suggest that other water sources, such as groundwater, may be involved in water formation in Lake Jiro.

**Fig 6.**
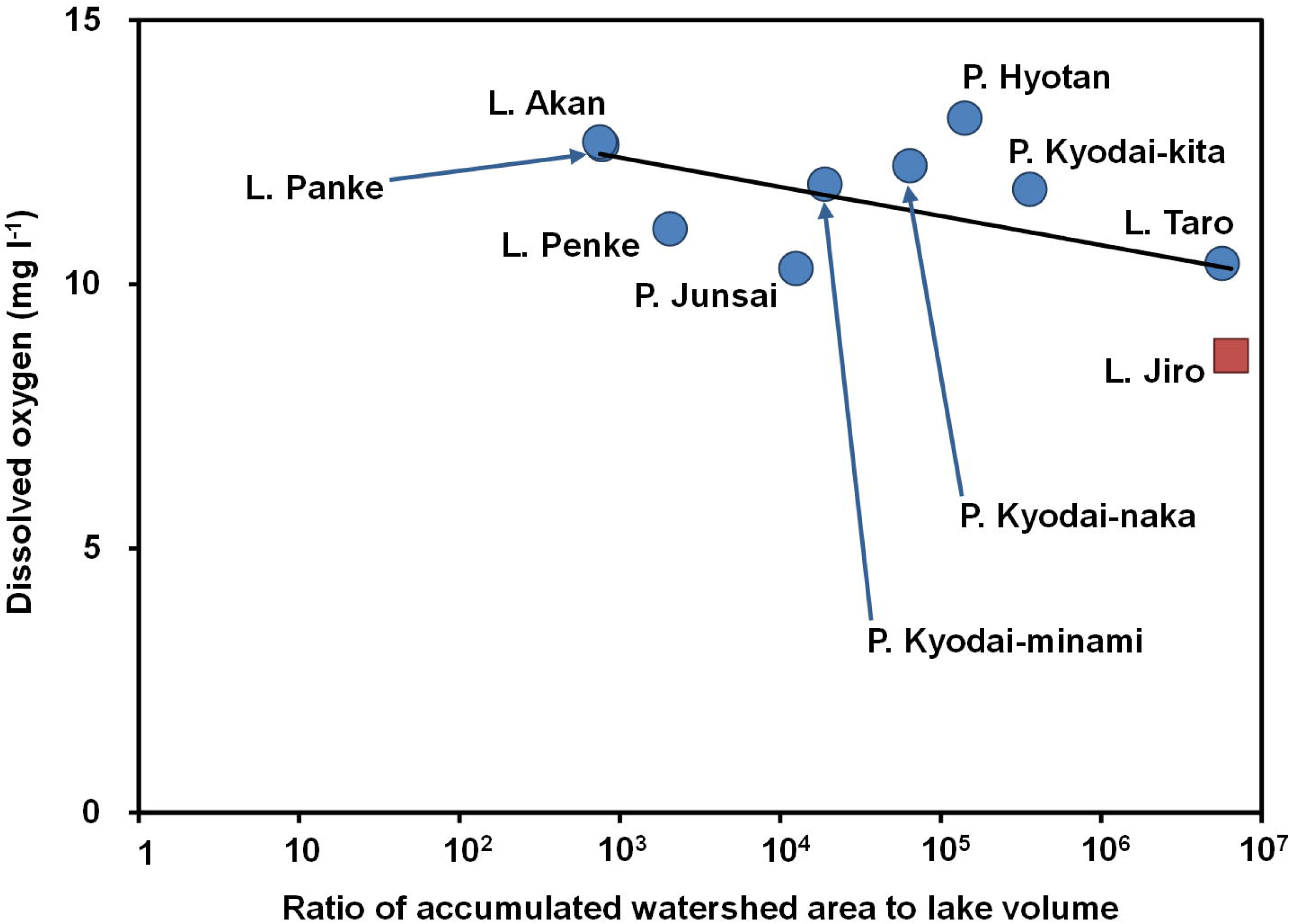
Relationship between ratios of accumulated watershed area to lake volume and dissolved oxygen (DO). Despite high correlation coefficients (Table 1), the regression line may not indicate an environmental gradient of DO, because its slight negative slope results from the uniquely low DO data for Lake Jiro (red square) with a relatively heterogeneous water quality (S1 Table).

Finally, regardless of the lake’s size, water temperature decreases with elevation upstream, and the difference in water temperature was 3.5 °C for an elevation difference of about 150 m (Fig 7).

**Fig 7.**
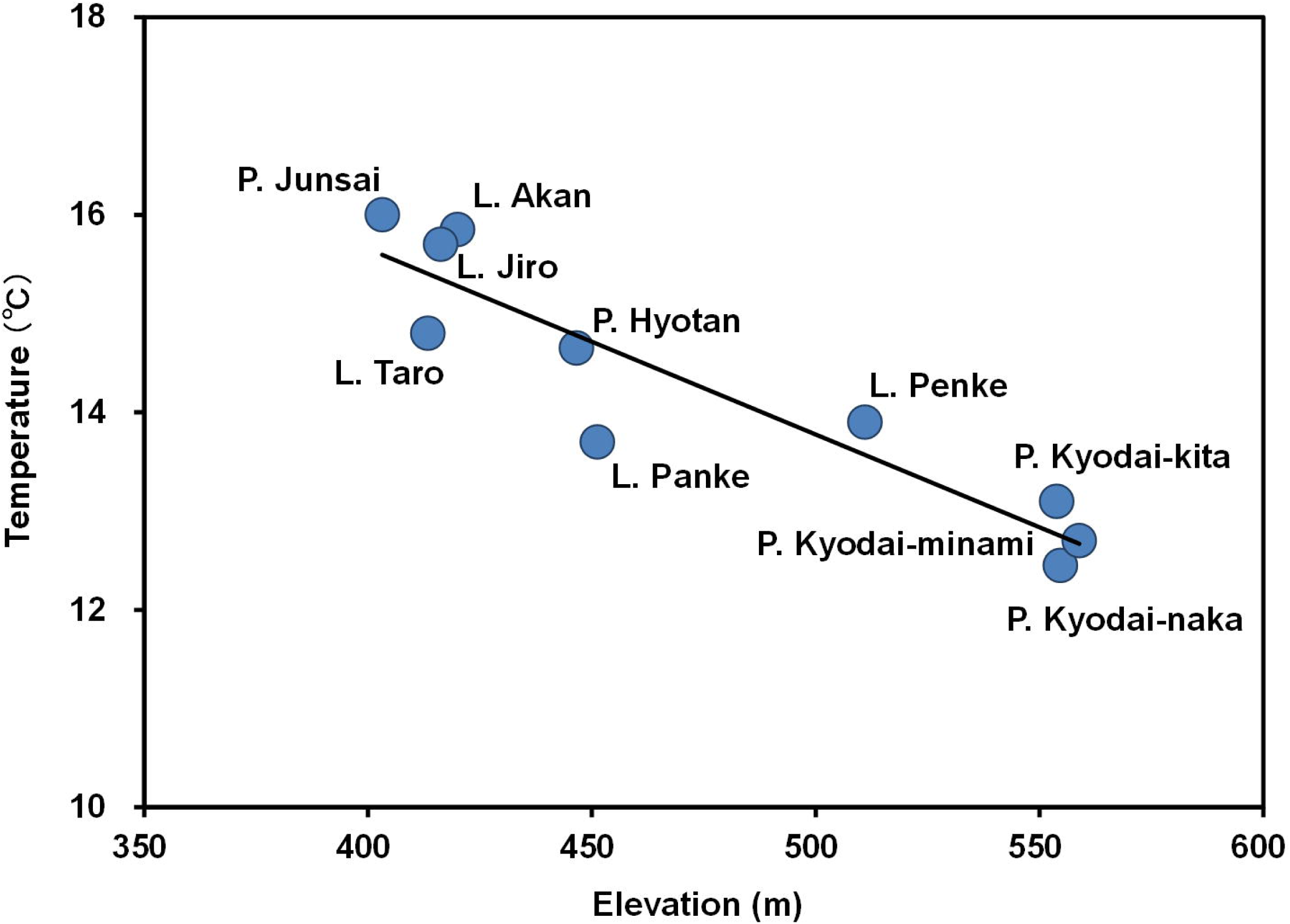
Relationship between elevation and water temperature in Akan Caldera Lakes. Water temperature is lower in lakes located upstream at higher elevations, regardless of lake size.

### Distribution and species composition of macrophytes

We recorded 21 species of macrophytes in total (excluding emergent plants and macro algae) in 7 lakes, while no macrophyte species were observed in 3 lakes without inflow rivers (S1 Table). The correlation matrix of the number of macrophyte species was strong and significant with the following variables among the above-mentioned lake topographic characters (S2 Table): boundary length (r = 0.720, *p* < 0.05), shoreline development (r = 0.703, *p* < 0.05), maximum depth (r = 0.924 < 0.001), mean depth (Fig 8, r = 0.928, *p* < 0.001) and residence time (r = 0.921, *p* <0.001). Several previous studies on relatively shallow or small lakes and ponds reported that the number of macrophyte species is correlated with lake area and that MacArthur and Wilson’s “theory of island biogeography”, which theorizes that larger islands are more biodiverse than smaller islands, is applicable here [52–59]. However, in our study no significant correlation was found between lake area and the number of macrophyte species (S2 Table). The boundary length and the shoreline development are parameters affecting horizontal length of the littoral zone, and the magnitude of the maximum and mean depths and the residence time related to depth and volume of lake basin contribute to expansion of the vertical littoral zone under conditions of greater water clarity. Therefore, the number of macrophyte species in the large lakes of Akan Caldera is presumed to be more closely related to the littoral zone area than the lake area.

**Fig 8.**
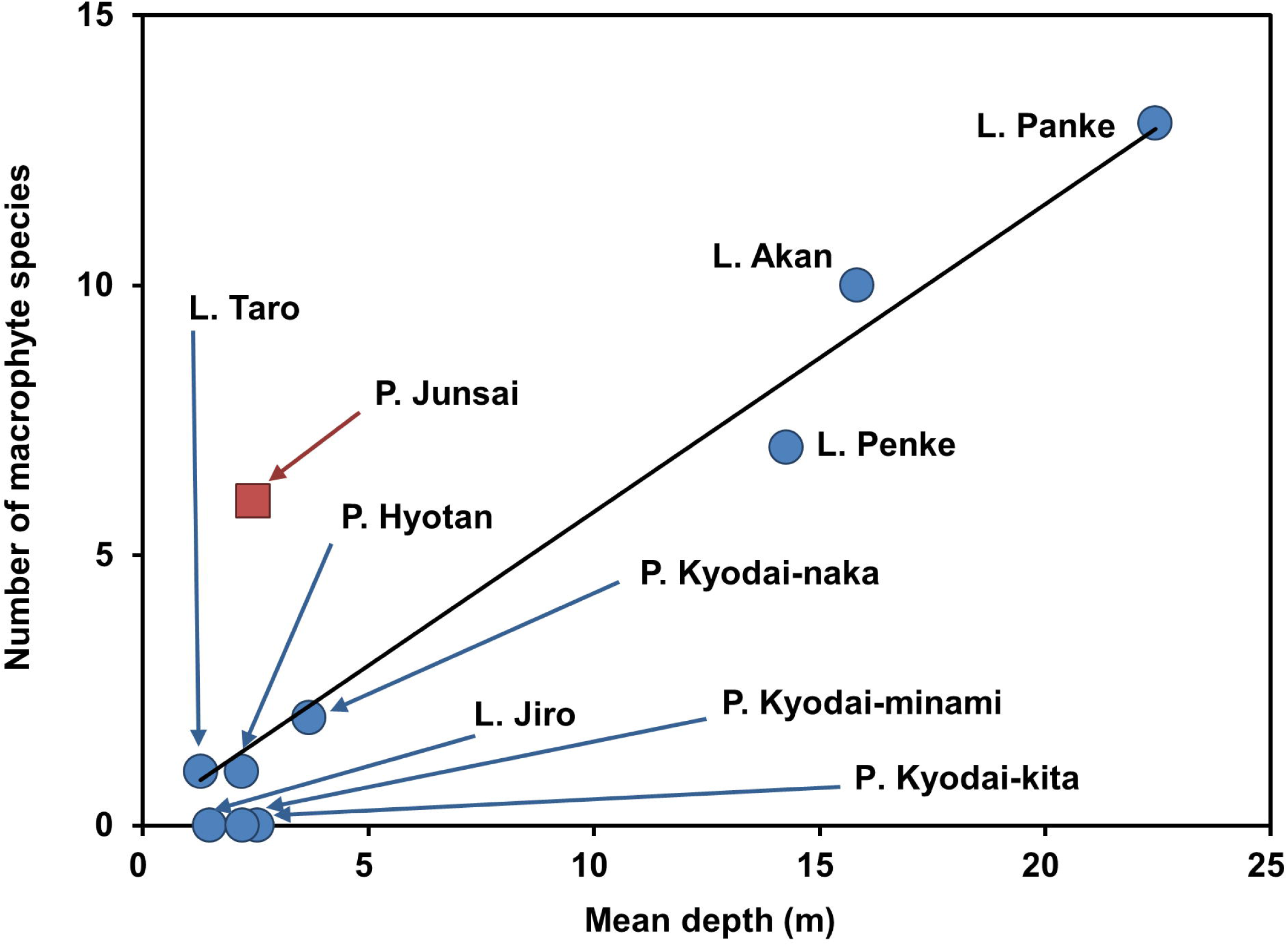
Effect of lake size on number of aquatic macrophyte species in Akan Caldera Lakes. In total, 21 species were observed in 7 lakes. The number of species in each lake shows high correlation with mean depth as well as with maximum depth, boundary length, shoreline development and residence time (S2 Table). Pond Junsai (red square) has dystrophic water quality (S1 Table) and is dominated by floating-leaved and free-floating plants, whereas most other lakes have a submerged plant community (S1 Table).

The two-dimensional plots of the number of macrophyte species and the above five topographic characters showed that Pond Junsai, a dystrophic lake, was designated an outlier (Fig 8). Furthermore, when the individual species was classified as submerged, floating-leaved and free-floating, the correlation coefficient was significantly greater with the number of species of submerged plants (S2 Table). The number of species of floating-leaved and free-floating plants showed significant and strong correlation with the water quality parameters Chl-a, DOC, COD and TN (S2 Table), with most of these species localized in Pond Junsai. Thus, we conducted a cluster analysis to understand species composition of macrophytes in each lake (Fig 9). The cluster was firstly divided into two groups: 6 species of floating-leaved and free-floating plants of Pond Junsai, and 14 submerged and 1 floating-leaved plants in the other lakes. Pond Junsai contained some indicator species of the dystrophic water: *Brasenia schreber*i (Fig 10), *Nuphar pumila* var. *pumila*, *Nymphaea tetragona* var. *tetragona* and *Utricularia macrorhiza* [60]. The remaining species recorded from the other lakes were classified as oligotrophic, mesotrophic, oligo-mesotrophic and oligo-meso-eutrophic, following the lake type based on the trophic status shown in Fig 2A. *Ranunculus nipponicus* var. *submersus* (Fig 10), *Potamogeton alpinus* and *Isoetes asiatica*, typically found in more pristine water [60–62], were designated oligotrophic. *Myriophyllum spicatum* (Fig 10), *Hydrilla verticillata*, *Potamogeton compressus, Potamogeton pectinatus* and *Ceratophyllum demersum,* occurring in more eutrophic settings [38, 60, 61], were designated oligo-mesotrophic. *Potamogeton crispus* (Fig 10), distributed in a variety of water environments [60], was designated mesotrophic.

**Fig 9.**
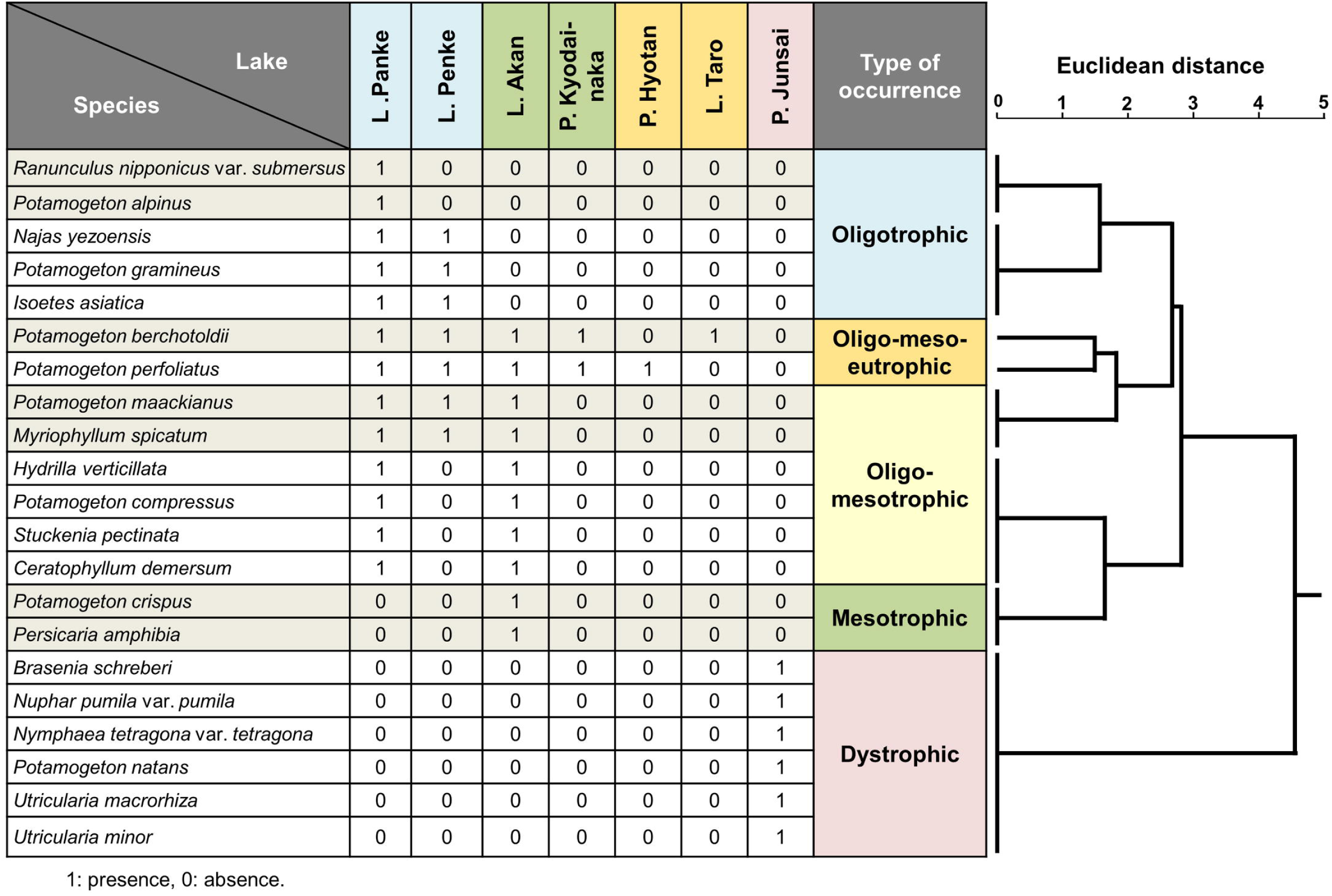
Cluster analysis of distribution and species composition of aquatic macrophytes in Akan Caldera Lakes. The tree diagram was drawn by the Ward method using Euclid distance based on the 1/0 data converted from the presence/absence data for the seven lakes where macrophytes occurred. According to the lake habitat types (Fig. 2A), clusters are classified into five vegetation types: oligotrophic, mesotrophic, oligo-mesotrophic, oligo-meso-eutrophic, and dystrophic.

**Fig 10.**
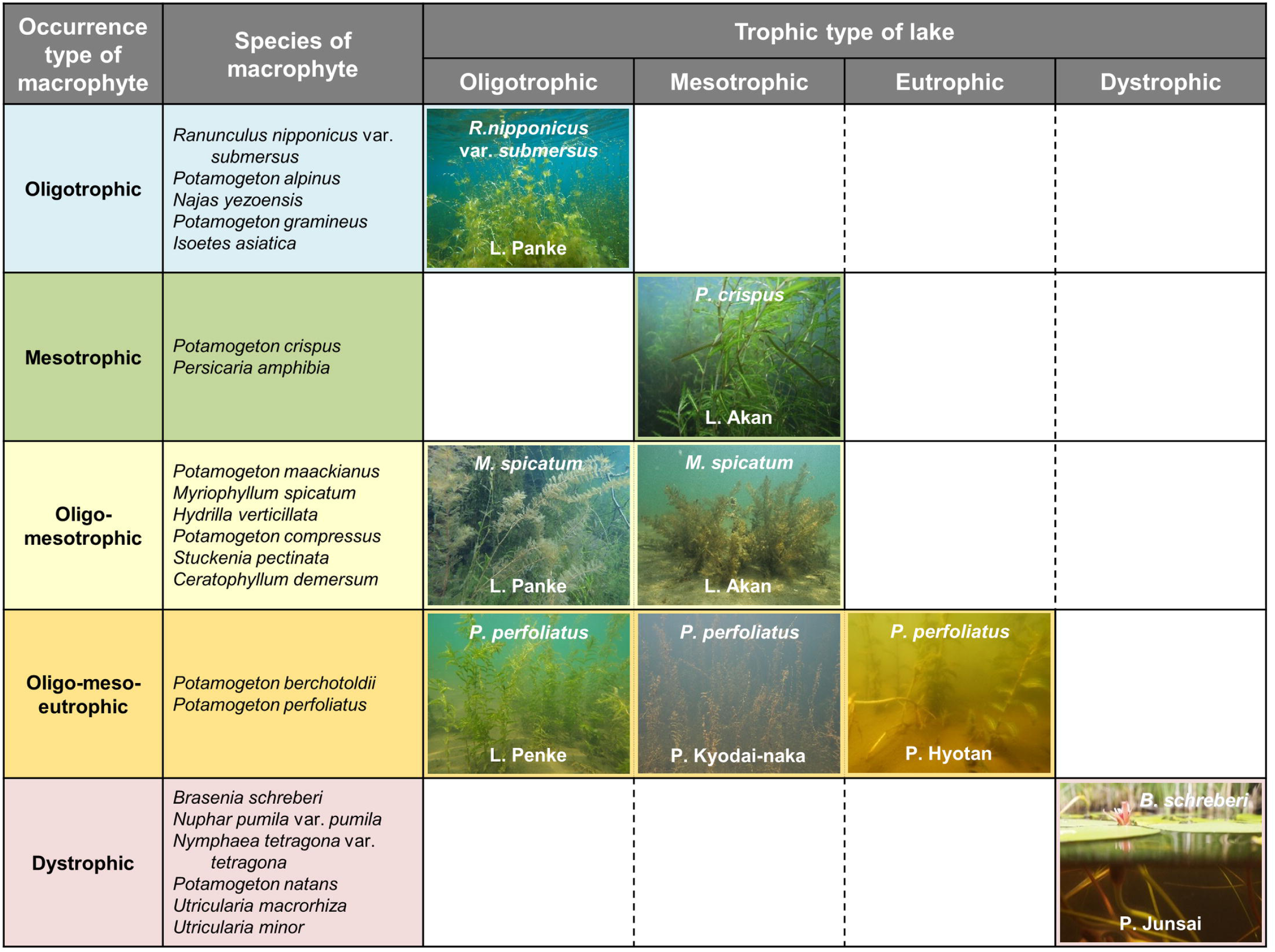
Determination model for macrophyte species composition corresponding to trophic types in Akan Caldera Lakes. Individual macrophyte species belong to any one of the five occurrence types of macrophytes, accompanied by inherent trophic type(s) and range. Macrophytes of oligotrophic-, mesotrophic- and dystrophic-occurrence types are only distributed within oligotrophic-, mesotrophic- and dystrophic-type lakes, respectively. In contrast, within oligo-mesotrophic- and oligo-meso-eutrophic-occurrence types, macrophytes are able to occur widely in oligotrophic/mesotrophic- and oligotrophic/mesotrophic/eutrophic-type lakes. Accordingly, species composition of macrophytes in oligotrophic- and mesotrophic-type lakes is determined by combination of the occurrence types, oligotrophic/oligo-mesotrophic/oligo-meso-eutrophic and mesotrophic/oligo-mesotrophic/oligo-meso-eutrophic. In eutrophic and dystrophic lakes, oligo-meso-eutrophic- and dystrophic-occurrence types are distributed, respectively.

As mentioned above, prior to anthropogenic eutrophication in the first half of the 20th century, the phosphorus level in Lake Akan appears to have been much lower than currently (Fig 3). Thus, the current aquatic flora was also compared with results from the oldest known vegetation survey conducted in 1897 [63]. Of the ten species of macrophytes in Lake Akan observed in our study, nine species were also found in 1897, except for *P. crispus,* classified as mesotrophic. However, the two oligotrophic species *R. nipponicus* var. *submersus* and *I. asiatica* in the old list were not found. These results suggest a shift to a more eutrophic vegetation type.

### Uniqueness of Akan Caldera Lakes

To summarize, although the lakes of Akan Caldera have the same origin, after being divided by a volcanic eruption they have developed into a series of oligotrophic, mesotrophic, eutrophic and dystrophic lakes (Fig 2A). The trophic level of each lake indicated by TP was closely related to the ratio of watershed area (AWA) to lake size (LA and LV) (Fig 2B). These results strongly suggest the possibility that the rate of eutrophication was different among the lakes, and we see the various stages of lake succession in progress in this system. However, the observed TP of Lake Akan and its downstream neighbors, Lake Jiro and Lake Taro, are assumed to have been impacted by anthropogenic eutrophication in the past (Fig 3A). Furthermore, the formation history of the Akan Caldera Lakes suggests that Ponds Hyotan and Junsai are significantly older than the other lakes [27–29]. As shown in Fig 2A, Chl-a levels in these two ponds are specifically higher than those of other lakes, which might be due to differences in time of formation. To understand the long-term changes in nutrient loading, including TP, and in primary production, the correct time of formation and subsequent eutrophication history of each lake should be clarified through research on lake sediment, etc. In addition, the linear regression of the relationship between TP and AWA/LA and AWA/LV (Fig 2B) suggests that the indigenous environmental variables of the watersheds may vary little. However, in reality, the geology and vegetation within the Akan Caldera are not uniform [27, 30], and further research is needed to determine the actual phosphorus loading from the watersheds.

Macrophyte species composition varied among lakes, driven by lake trophic conditions (Fig 9), and is indicative of aquatic plant succession, or “hydrarch succession”. Many species of macrophytes are classified into a variety of “types” according to environmental characteristics of habitats, including trophic level [56, 61, 64–68]. Schneider and Melzer [68], for example, proposed seven categories ranging from oligotrophic to eutrophic and even polytrophic types. Meanwhile, Lacoul and Freedman [61] simplified into three categories: oligotrophic, eutrophic, and general types, based on the opinion that many macrophytes generally have a broad ecological range, occurring over wide trophic levels, while other species have a narrower distribution. In our study, however, all of the observed species belonged to only one of the five occurrence types, with a species-specific range of trophic level (Fig 10). Thirteen species were distributed among oligotrophic, mesotrophic and dystrophic lake types, respectively, and eight species occurred in the oligotrophic and mesotrophic, and the oligotrophic, mesotrophic and eutrophic lakes with wider trophic ranges. These results explain how the species composition of macrophytes in the Akan Caldera Lakes with different trophic types is determined by the combination of species with different trophic requirements. Importantly, we noted vegetation changes in Lake Akan due to anthropogenic eutrophication: this was characterized by disappearance of oligotrophic-type species, appearance of mesotrophic-type species, and survival of oligo-mesotrophic and oligo-meso-eutrophic-type species, when the trophic level of Lake Akan shifted from oligotrophic to mesotrophic as shown in Fig 10. This might be the first example to clearly and simply illustrate the replacement of species in an aquatic plant community caused by trophic level change. Further investigation is necessary to test whether this is a universal phenomenon in limnology.

Trophic levels are not the only factor known to affect macrophyte distribution and species composition. At a smaller scale, the following physical factors have been important in producing environmental gradients between or within lakes: topography, geological qualities, inflow waters as physical factors in the watershed, lake basin morphology (mainly depth and area), water temperature, light conditions, turbidity, current flow, substrate (sediment). And, chemical factors in play include inorganic ions, salinity, organic matter, conductivity, alkalinity, pH and nutrients [11, 13, 21, 54, 56, 59, 61, 62, 64, 65, 69–73]. As this itemization indicates, a comprehensive understanding of the mechanisms that determine the distribution and species composition of macrophytes is difficult, due to multiple influencing factors. In the Akan Caldera Lakes, species distribution was related to trophic conditions (Fig 9 and 10), while number of species was closely related to lake size (Fig 8). In general, species diversity at smaller scales increased depending upon spatial environmental heterogeneity [74, 75]. Lakes Akan, Panke and Penke, all large and oligo- and mesotrophic lakes with high species count, appear to offer a large variety of the physical and chemical factors noted above. The topography of capes, bays and islands in these lakes diversifies wave action and substrate via varying wind-wave parameters, and “fetch”, i.e. the length of the lake surface over which wind blows [4, 61]. Inflow rivers locally alter substrate, current and water quality, greater water depth lowers water temperature and reduces substrate grain size, and oligo- or mesotrophic water allows sunlight to penetrate deeper into the littoral zone, leading to a gradual gradient in the light environment [4, 5, 21, 55, 56, 59, 61, 62, 69]. This environmental variability may offer habitat for a number of macrophytes in these large lakes. On the other hand, Pond Junsai, dominated by floating-leaved and free-floating plants, is the only example of dystrophic water quality in the Akan Caldera Lakes. Its brownish lake water, derived from high DOC containing abundant humic substances, suppresses submerged plant growth due to high light absorption [5, 21, 56, 61, 62]. Humic substances are thought to originate from decomposing terrestrial and/or littoral plant material [5, 22, 50, 61]. Thus, eutrophication in Pond Junsai may have undergone a different process than in the other lakes because of differing surrounding vegetation, even if the initial process was the same. To understand the factors affecting plant distribution the relationships between detailed macrophyte distribution and habitat micro-environments in each lake, including impacts of the surrounding vegetation, must be elucidated.

## Conclusions

In this study, we demonstrated that the ratio of Akan Caldera watershed area to lake size was positively correlated with TP concentration, an indicator of trophic level in lakes. Furthermore, the variety of trophic levels in the lakes, ranging from oligotrophic to eutrophic, coincided with the “lake succession” or “lake type” series of ecological succession in lakes. Further, “hydrarch succession”, a change in the species composition of macrophytes, also indicates a regular series corresponding to the trophic levels. These facts indicate that the unique eutrophication rates, defined by the ratio of watershed area to lake size, may have diversified the water quality and the aquatic flora of the lakes. Uniquely, this possibly illustrates lake ecological succession in “real time”, previously thought impossible to see [2, 9, 18, 24]. In other words, the Akan Caldera Lakes can be seen as a massive experiment conducted by nature in an “Akan Caldera Laboratory”, or seen as a showcase visualizing a freshwater ecosystem, focusing on lake and hydrarch succession. The essence of ecological succession is, simply put, nothing more than a developing and changing process of biodiversity [75]. In this context, the Akan Caldera Lakes will enable a new approach to the comparative study of spatio-temporal fluctuations in biodiversity and aquatic environments, including responses to changes in the surrounding environment, especially climate change, now seen as an issue of paramount importance [24, 26].

## Acknowledgements

A part of this study is “Research of the aquatic plants in Lake Akan and circumference lakes” and “Lake charts drawing of Lake Akan and circumference lakes” conducted in 2013 and 2014 by the Ministry of Environment of Japan. The City of Kushiro provided facilities and funding for compilation of the study. Árni Einarrson, Mývatn Research Station, Iceland, provided advice on research design. Takehiko Fukushima, University of Tsukuba and Koichi Kamiya, Ibaraki Kasumigaura Environmental Science Center supported the water analyses. The Hokkaido Shimbun Press contributed to the aerial photography. Satoshi Kofuku and Kokoro kikuchi, IDEA Consultants, Inc., assisted with literature research. Shuji Hino, Yamagata University and Yasushi Ishikawa, Hokkaido Research Organization, advised regarding caldera lakes and calculation of water quality. Mami Yamazaki, Sapporo Museum Activity Center, provided advice on macrophytes.

## Supporting information

S1 Table. Data set of topography, water quality and number of species of macrophytes in Akan Caldera Lakes.

S2 Table. Correlation matrix of topography, water quality and number of species of macrophytes in Akan Caldera Lakes.

## Notes

### Competing Interest Statement

The authors have declared no competing interest.

## References

1. Whittaker RH. Communities and ecosystems. 2nd ed. New York: Macmillan; 1975.

2. Sakamoto M. Ecological succession, II. Tokyo: Kyoritsu-Shuppan; 1976. (in Japanese)

3. Owen OS. Natural resource conservation: an ecological approach, 4th ed. New York: Macmillan; 1985.

4. Horne AJ, Goldman CR. Limnology, 2nd ed. New York: McGraw-Hill; 1994.

5. Wetzel RG. Limnology: lake and river ecosystems, 3rd ed. San Diego: Academic Press; 2001.

6. Schindler DW, Vallentyne JR. The algal bowl, overfertilization of the world’s freshwaters and estuaries. Edmonton: University of Alberta Press; 2008.

7. Vollenweider R, Kerekes J. Eutrophication of waters: Monitoring, assessment and control. Paris: Organization for Economic Co-Operation and Development (OECD); 1982.

8. Lindeman RL. The trophic-dynamic aspect of ecology. Ecology. 1942;23: 399–417. https://doi.org/10.2307/1930126.

9. Forel FA. Handbuch der Seenkunde: allgemeine Limnologie. Stuttgart: J. Engelhorn; 1901. (in Germany)

10. Yoshimura S. Limnology. Tokyo: Sanseido; 1937. (in Japanese)

11. Pearsall WH. The development of vegetation in the English lakes, considered in relation to the general evolution of glacial lakes and rock basins. Proc R Soc B, Biol Sci. 1921;92: 259–284. https://doi.org/10.1098/rspb.1921.0024.

12. Dachnowski A. Peat deposits of Ohio: their origin, formation and uses. Columbus: Division of Geological Survey, Ohio; 1912. http://hdl.handle.net/1811/78409

13. Pearsall WH. The aquatic vegetation of the English lakes. J. Ecol. 1920;8: 163–201. doi: 10.2307/2255612.

14. Dachnowski AP. Factors and problems in the selection of peat lands for different uses. Washington, D. C.: United States Department of Agriculture; 1926. https://doi.org/10.5962/bhl.title.108906

15. Kormondy EJ. Concepts of ecology. 2nd ed. Englewood Cliffs: Prentice-Hall; 1969. http://ci.nii.ac.jp/ncid/BA1127514X

16. Odum EP. Fundamentals of ecology, 3rd ed. Philadelphia: Saunders; 1971.

17. Morishita I. Dam lake ecology. Tokyo: Sankaidou; 1983. (in Japanese)

18. Margalef R. Perspectives in ecological theory. Chicago: University of Chicago Press; 1968.

19. Odum EP. The strategy of ecosystem development. Science. 1969;164: 262–270. http://habitat.aq.upm.es/boletin/n26/aeodu.en.html

20. Wetzel RG, Likens GE. Limnological analyses, 3ed ed. New York: Springer; 2000.

21. Dodson SI. Introduction to limnology. New York: McGraw-Hill; 2005.

22. Dodds WK, Whiles MR. Freshwater ecology, 2nd ed. Burlington: Academic Press; 2010.

23. Smith J, Appleby P, Battarbee R, Dearing J, Flower R, Haworth E, et al. Environmental history and palaeolimnology. Springer Science & Business Media; 1991.

24. Kling GW. A lake’s life is not its own. Nature. 2000;408: 149–150. doi: https://doi.org/10.1038/35041659.

25. Johnson EA, Miyanishi K. Testing the assumptions of chronosequences in succession. Ecol Lett. 2008;11: 419–431. doi: https://doi.org/10.1111/j.1461-0248.2008.01173.x

26. Sayer C, Roberts N, Sadler J, David C, Wade P. Biodiversity changes in a shallow lake ecosystem: a multi - proxy palaeolimnological analysis. J Biogeogr. 1999;26: 97–114. doi: https://doi.org/10.1111/j.1365-2699.1999.00298.x

27. Satoh H. Akanko, explanatory text of the geological map of Japan, Kushiro, No. 7. Tokyo: Geological Survey of Japan; 1965. (in Japanese with English summary)

28. Tamada J, Nakagawa M. Eruption history of Oakan volcano, eastern Hokkaido, Japan. Kazan. 2009;54: 147–162. (in Japanese with English summary)

29. Hasegawa T, Nakagawa M. Large scale explosive eruptions of Akan volcano, eastern Hokkaido, Japan: a geological and petrological case study for establishing tephro-stratigraphy and-chronology around a caldera cluster. Quat Int. 2016;397: 39–51. doi: https://doi.org/10.1016/j.quaint.2015.07.058.

30. Maeda Ippoen Foundation. The nature of Akan National Park, 1993. Akan: Maeda Ippoen Foundation; 1994. (in Japanese)

31. Hino S, Mikami H, Arisue J, Ishikawa Y, Imada K, Yasutomi R, et al. Limnological characteristics and vertical distribution of phytoplankton in oligotrophic lake Akan-Panke. Jpn J Limnol. 1998;59: 263–279. doi: https://doi.org/10.3739/rikusui.59.263

32. Tanaka M. The lakes in Japan. Nagoya: Nagoya University Press; 1992. (in Japanese)

33. Hokkaido Institute of Environmental Science. Lakes and marshes in Hokkaido, revised edition. Sapporo: Hokkaido Institute of Environmental Science; 2005. (in Japanese)

34. Clements FE. Plant succession: an analysis of the development of vegetation. Washington: Carnegie Institution of Washington; 1916. http://ci.nii.ac.jp/ncid/BA39490309

35. Solińska-Górnicka B, Symonides E. Long-term changes in the flora and vegetation of Lake Mikołajskie (Poland) as a result of its eutrophication. Acta Soc Bot Pol. 2001;70: 323–334. doi: https://doi.org/10.5586/asbp.2001.040.

36. Kowalczewski A, Ozimek T. Further long-term changes in the submerged macrophyte vegetation of the eutrophic Lake Mikolajskie (North Poland). Aquat Bot. 1993;46: 341–345. doi: https://doi.org/10.1016/0304-3770(93)90013-M.

37. Coops H, Doef RW. Submerged vegetation development in two shallow, eutrophic lakes. Hydrobiol. 1996;340:115–120. doi: https://doi.org/10.1007/BF00012742.

38. Egertson CJ, Kopaska JA, Downing JA. A century of change in macrophyte abundance and composition in response to agricultural eutrophication. Hydrobiol. 2004;524:145–156. doi: https://doi.org/10.1023/B:HYDR.0000036129.40386.ce.

39. Tanakadate S. Research report on the volcano lakes in Hokkaido. Sapporo: Hokkaido Government; 1925. (in Japanese)

40. Takayasu S, Igarashi H, Sawa K. Lake survey, Lake Akan. Sci Rep Hokkaido Fish Exp Stn. 1930;21: 67–92. (in Japanese)

41. Takayasu S, Kondo K. River and lake survey, Lake Panke and Lake Penke. Sci Rep Hokkaido Fish Exp Stn. 1936;39: 1–24. (in Japanese)

42. Sugimoto T. Introduction to Kashmir C3. Tokyo: Jitsugyo no Nihon Sha; 2010. (in Japanese)

43. Umekawa T, Wakana I, Ohara M. Reproductive behavior and role in maintaining an aggregative form of the freshwater green alga Marimo, *Aegagropila linnaei*, in Lake Akan, Hokkaido, Japan. Aquat Bot. 2021;168: 103309. doi: https://doi.org/10.1016/j.aquabot.2020.103309.

44. Igarashi S, Ishikawa Y, Mikami H. Long-term changes in limnological characteristics in Lake Akan, Hokkaido. Res Rep NIES, Japan. 2000;153: 34–54. (in Japanese)

45. K0102 J. Testing Methods for Industrial Wastewater. Tokyo: Japanese Standards Association; 2013. (in Japanese)

46. Grasshoff K, Kremling K, Ehrhardt M. Methods of seawater analysis: Weinheim: John Wiley & Sons; 2009.

47. SCOR-UNESCO. Determination of photosynthetic pigments in seawater. Monographs on oceanographic methodology 1. New York: UNESCO Publications Center; 1966.

48. Fukushima T, Imai A, Matsushige K, Aizaki M, Otsuki A. Freshwater DOC measurements by high-temperature combustion: comparison of differential (DTC-DIC) and DIC purging methods. Water Res. 1996;30: 2717–2722. doi: https://doi.org/10.1016/S0043-1354(96)00198-4.

49. Miyadi D. Studies on the bottom fauna of Japanese lakes, VII. Lakes of Hokkaido. Jpn J Zool. 1932;4: 223–252.

50. Brönmark C, Hansson L-A. The biology of lakes and ponds, 2nd ed. Oxford: Oxford University Press; 2005.

51. Motoda S. The lakes in Hokkaido Sci Rep Hokkaido Fish Hatchery. 1950;5: 1–96. (in Japanese)

52. Wilson EO, MacArthur RH. The theory of island biogeography. Princeton: Princeton University Press; 1967.

53. Møller TR, Rørdam CP. Species numbers of vascular plants in relation to area, isolation and age of ponds in Denmark. Oikos. 1985;45: 8–16. doi: https://doi.org/10.2307/3565216.

54. Rørslett B. Principal determinants of aquatic macrophyte richness in northern European lakes. Aquat Bot. 1991;39: 173–193. doi: https://doi.org/10.1016/0304-3770(91)90031-Y.

55. Weiher E, Boylen CW. Patterns and prediction of α and β diversity of aquatic plants in Adirondack (New York) lakes. Can. J. Bot. 1994;72: 1797–1804. doi: doi.org/10.1139/b94-221.

56. Toivonen H, Huttunen P. Aquatic macrophytes and ecological gradients in 57 small lakes in southern Finland. Aquat Bot. 1995;51: 197–221. doi: https://doi.org/10.1016/0304-3770(95)00458-C.

57. Linton S, Goulder R. Botanical conservation value related to origin and management of ponds. Aquat Conserv Mar Freshw Ecosyst. 2000;10: 77–91. doi: https://doi.org/10.1002/(SICI)1099-0755(200003/04)10:2<77::AID-AQC391>3.0.CO;2-Y.

58. Mäkelä S, Huitu E, Arvola L. Spatial patterns in aquatic vegetation composition and environmental covariates along chains of lakes in the Kokemäenjoki watershed (S. Finland). Aquat Bot. 2004;80: 253–269. doi: https://doi.org/10.1016/j.aquabot.2004.08.006.

59. Søndergaard M, Jeppesen E, Jensen JP. Pond or lake: does it make any difference? Arch Hydrobiol. 2005;162: 143–165. doi: 10.1127/0003-9136/2005/0162-0143.

60. Kadono Y. A field guide to aquatic plants of Japan. Tokyo: Bun-ichi Sogo Shuppan; 2014. (in Japanese)

61. Lacoul P, Freedman B. Environmental influences on aquatic plants in freshwater ecosystems. Environ Rev. 2006;14: 89–136. doi: https://doi.org/10.1139/a06-001.

62. Søndergaard M, Johansson LS, Lauridsen TL, Jørgensen TB, Liboriussen L, Jeppesen E. Submerged macrophytes as indicators of the ecological quality of lakes. Freshw Biol. 2010;55: 893–908. doi: https://doi.org/10.1111/j.1365-2427.2009.02331.x.

63. Kawakami T. Collection record in Akan, Kushiro Province. Bot Mag Tokyo. 1898;12: 220–225. (in Japanese)

64. Seddon B. Aquatic macrophytes as limnological indicators. Freshw Biol. 1972;2: 107–130. doi: doi.org/10.1111/j.1365-2427.1972.tb00365.x.

65. Wiegleb G. A study of habitat conditions of the macrophytic vegetation in selected river systems in Western Lower Saxony (Federal Republic of Germany). Aquat Bot. 1984;18: 313–352. doi: https://doi.org/10.1016/0304-3770(84)90055-X.

66. Mäkirinta U. Classification of South Swedish Isoetid vegetation with the help of numerical methods. Vegetatio. 1989;81: 145–157. https://link.springer.com/chapter/10.1007/978-94-009-2432-1_12.

67. Heegaard E, Birks HH, Gibson CE, Smith SJ, Wolfe-Murphy S. Species–environmental relationships of aquatic macrophytes in Northern Ireland. Aquat Bot. 2001;70: 175–223. doi: https://doi.org/10.1016/S0304-3770(01)00161-9.

68. Schneider S, Melzer A. The trophic index of macrophytes (TIM)–a new tool for indicating the trophic state of running waters. Internat Rev. Hydrobiol. 2003;88: 49–67. doi: https://doi.org/10.1002/iroh.200390005.

69. Hutchinson GE. A treatise on limnology, Vol III, Limnological Botany. New York: John Wiley; 1975.

70. Spence DHN. Factors controlling the distribution of freshwater macrophytes with particular reference to the lochs of Scotland. J Ecol. 1967;55: 147–170. doi: 10.2307/2257723.

71. Kadono Y. Occurrence of aquatic macrophytes in relation to pH, alkalinity, Ca^++^, Cl and conductivity. Jap J Ecoi. 1982;32: 39–44. doi: 10.18960/seitai.32.1_39.

72. Vestergaard O, Sand-Jensen K. Alkalinity and trophic state regulate aquatic plant distribution in Danish lakes. Aquat Bot. 2000;67: 85–107. doi: https://doi.org/10.1016/S0304-3770(00)00086-3.

73. Lukács BA, Tóthmérész B, Borics G, Várbíró G, Juhász P, Kiss B, et al. Macrophyte diversity of lakes in the Pannon Ecoregion (Hungary). Limnologica. 2015;53: 74–83. doi

74. Wright DH, Currie DJ, Maurer BA. Energy supply and patterns of species richness on local and regional scales. In: Ricklefs RE, Schluter D, editors. Species diversity in ecological communities: historical and geographical perspectives. 7. Chicago: University of Chicago Press; 1993.

75. Miyashita T, Noda T. Community ecology. Tokyo: University of Tokyo Press; 2003. (in Japanese)

